# Quality circles for quality improvement in primary health care: their effectiveness, gaps of knowledge, origins and significance – a scoping review

**DOI:** 10.1101/387605

**Authors:** Adrian Rohrbasser, Janet Harris, Sharon Mickan, Geoff Wong

**Author notes:** **Corresponding author:** General Practitioner Medbase; Friedtalweg 18, 9500 Wil / Switzerland.

## Abstract

**Background:** Quality circles, or similarly structured small groups in primary health care, such as peer review groups, consist of 6 to 12 professionals from the same background who meet regularly to improve their standard practice. This paper reports the results from a scoping search performed to clarify possible effectiveness, knowledge gaps, underlying concepts and significance.

**Objectives:** To gain insight into knowledge gaps and understanding of the effectiveness, origins and significance of quality circles.

**Methods:** A search strategy was developed starting with ‘quality circle’ in PubMed and the index terms from those articles revealed were then used as search terms to identify further papers. Repeating this process in collaboration with a librarian, search strings relating to quality circles were built, and databases searched up to December 2017. Any paper on structured quality circles or related small group work in primary health care was included when relevant to the objectives.

**Results:** From 11973 citations, 82 background papers and 58 key papers were identified, in addition to 12 books and 10 websites. 19 studies, one paper summarizing three studies and one systematic review suggest that quality circles can be effective in behaviour change, though with varying effect sizes. Quality circles and their techniques are complex, as they are not standardized, and changes seem to depend on the topic and context, which requires further research into how and why they work in order to improve them. From their origins in industry, they are now used in primary health care in many countries for continuous medical education, continuous professional development and quality improvement.

**Conclusion:** The evidence on quality circles indicates that they can successfully change general practitioner behaviour. As they are a complex intervention, theory-driven research approaches are needed to understand and improve their effectiveness. This is of major importance because they play an important role in quality improvement in primary health care in many countries.

## Background

Quality circles (QCs), also known as peer review groups, and other structured small groups that exist across Europe, are small groups of health care professionals who meet to reflect and improve their standard practice. They use various didactic methods, such as brain-storming and reflective thinking, and tools for quality improvement (QI), such as audit and feedback or purposeful use of local experts. They are used for quality initiatives in primary health care (PHC) in several European countries (1–10). Scotland and Wales recently introduced structured small groups for QI to replace a pre-existing outcomes-driven incentive scheme (11, 12). It is increasingly being recognized that what is missing from the literature is an account of effectiveness; namely, whether participants change their behaviour or not.

There are systematic reviews (SRs) on the tools used in these groups but there is still doubt as to whether they make participants improve their practice, even if the tools are used in combination. This paper reports the results of a scoping review to map areas of uncertainty regarding QC effectiveness, thereby indicating where further research is needed. It maps size and type of evidence and describes original intentions and reported benefits. To gain additional insight into the potential and significance in different countries, the historical development and drivers of QCs are also reported. The objectives of the scoping review involved several steps and followed the guidelines for conducting systematic scoping reviews (13):

- mapping the size and type of evidence of the existing literature
- describing and defining QCs
- establishing their effectiveness
- recognizing gaps in knowledge
- describing their intentions and reported benefits
- describing their historical development and significance

This paper provides a working definition of QCs and describes their basic properties, effectiveness, knowledge gaps, historical background and significance. The implications of possible knowledge gaps are also discussed.

## Methods

### Information Sources and Search

Background information, such as basic QC characteristics, reports on their historical development and their spread from industry to the health care sector, was retrieved from 12 relevant textbooks identified by AR, a content expert in the field of QI and associated small group work (14–25).

The literature search, including literature up to December 2017, was performed in several steps by AR. Initially, a limited search was performed in PubMed using the term ‘quality circle’ to identify some papers to be used as a starting point. In collaboration with an experienced librarian, analysis of text words in the title, abstract and indexing helped identify additional search terms. Iterative searching yielded search strings relating to descriptors of QCs, such as ‘quality improvement’, ‘group functions’ or ‘primary care’ (supplementary file S1). Literature was retrieved in Medline, Embase, PsycInfo and CINAHL without language or time restrictions and downloaded to Endnote, a standard software tool for publishing and managing bibliographies, citations and references.

### Eligibility Criteria

Any paper on QCs within PHC, with qualitative or quantitative outcomes, or background information, was considered for inclusion. AR screened all papers found during the search process, whilst SM, JH and GW cross-checked them for consistency in the application of the eligibility criteria.

### Paper Selection

The quality of data retrieved was only assessed as to whether it provided relevant information about QCs in PHC (26). AR made the relevance assessments, which were then explored in discussions with SM, JH and GW. The following questions were used to assess relevance:

- Does the paper cover the background of QCs in PHC?
- Does the paper describe the process in these small groups?
- Do the papers provide enough data to allow evaluation?

The number of papers excluded and included at each stage is indicated in the flow diagram (Figure 1).

**Figure 1:**
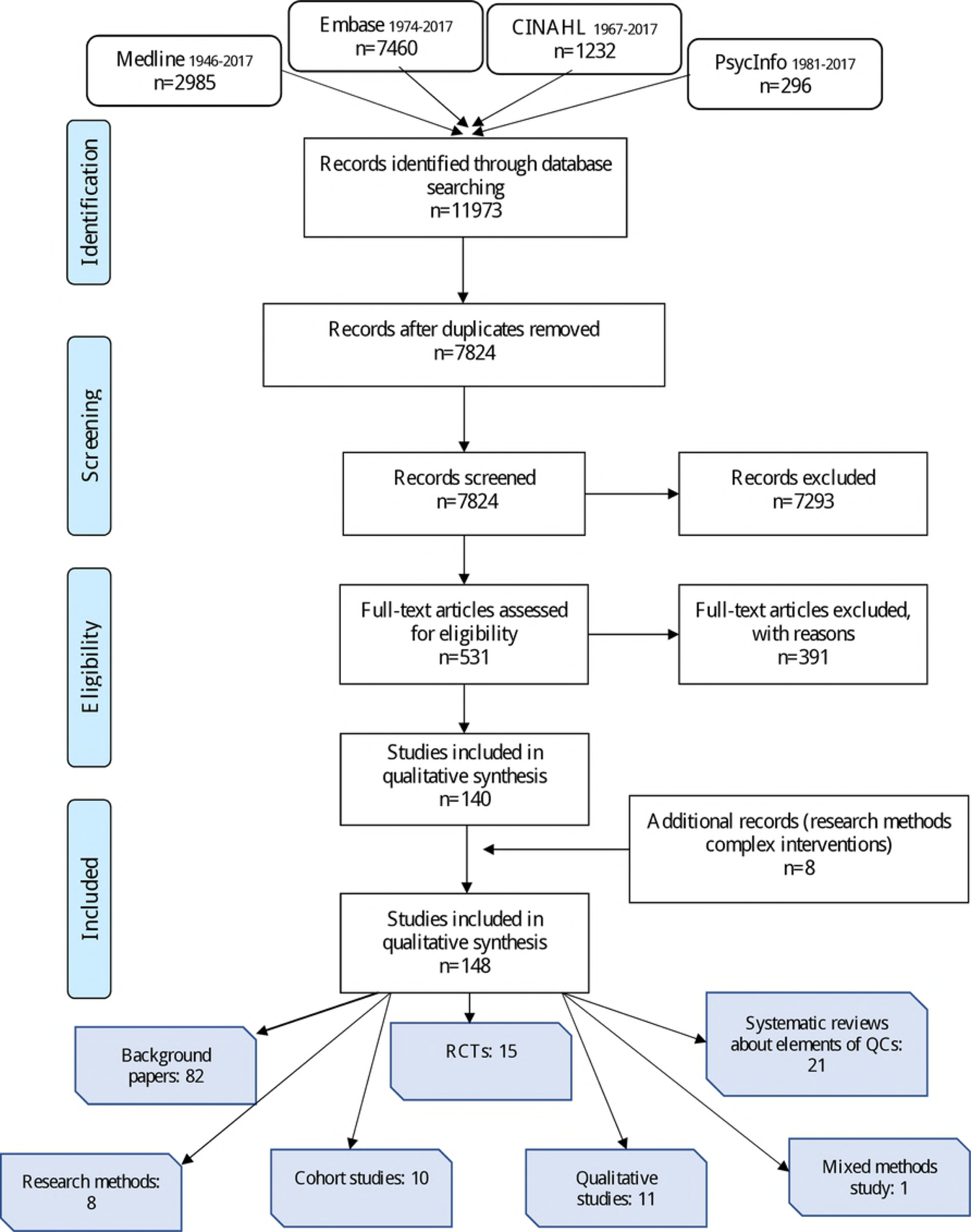
paper flow diagram

### Data Collection and Reporting

The following types of data were extracted from the documents by AR: authors, year of publication, location, and data describing background, definitions of QCs, their underlying processes and possible effectiveness, historical development and significance in PHC today. The data were then put into a narrative and tables to describe the different aspects of QCs. A data collection template was not used as it was difficult to anticipate how the data would be presented. Data were charted to answer the review objectives.

## Results

12 text books were identified, and iterative searches returned 82 background papers and 58 key papers which were deemed eligible and relevant (supplementary files S2, S3 and S4). Additionally, 10 informative websites of various organisations were identified (1–10) as well as 8 papers, after a specific search concerning the research methods of complex interventions, such as small group interventions (27–34). Key papers mainly described or evaluated processes of QCs using research methods, such as systematic reviews (SR), randomised controlled trials (RCT), cohort, or before and after studies. Qualitative studies provided further information on their process and additional benefits. Background papers and the aforementioned websites provided data in reports and summaries on the history, development and significance of QCs.

### What QCs are

Within those documents included, the authors identified concurrent key concepts relating to QCs and then agreed on a definition: QCs comprise small groups of 6 to 12 professionals from the same background who meet regularly to reflect on and improve their standard practice (1, 4–6, 8–10, 14, 16–23, 25, 35–46). The terms Practice Based Small Group Work, Peer Review Group, Problem Based Small Group Learning, Practice Based Research Group, Quality Circle, Continuous Medical Education (CME) Group, and Continuous Professional Development (CPD) Group are used interchangeably in different countries. The labelling suggests the basic, original intention of the group, although they may now serve the same purpose. In this scoping review, the term Quality Circle is used as an umbrella term to include all of them.

The groups choose a topic they want to learn more about or a quality aspect which they want to improve in their practice. They decide on how to approach and solve the issue, and they create space for reflective thinking to improve clinical practice (1, 5, 15, 22, 45, 47–55).

The groups also choose their own facilitators, who observe and lead the group through the cycle of QI. Whilst respecting the contribution of each individual, and taking into consideration group dynamics, facilitators try to keep the members focused on the issue without controlling them (19, 22, 56–61).

QC techniques usually comprise a combination of different types of tools, such as the use of educational material discussed in a workshop-like atmosphere, contact with local knowledge experts, audit and feedback on clinical practice with or without outreach visits, facilitation and local consensus processes (36, 37, 42, 43, 46, 54, 55, 62–67). The group may also rehearse clinical skills and use active didactic methods to promote learning, such as brain-storming, reflective thinking, self-monitoring and professional reprocessing of patient situations (1, 7, 8, 10, 18, 23, 25, 68).

The varying tools and didactic methods are usually tailored to the locally prevailing circumstances (30, 54, 68, 69). The number and difficulty of these tools and didactic methods, as well as outcomes and the context of the group, affect the process. Therefore, QCs are complex social interventions (28, 31, 70) that are run in PHC systems and which change constantly with the prevailing economic situation, scientific development and cultural circumstances (27, 30). They incorporate social aspects of the workplace that affect team work, self-determination and involvement in management at a day-to-day level.

### Effectiveness

24 quantitative studies and 1 SR were assessed as to whether QCs promote behaviour change. 19 studies, one paper summarizing three studies and one SR from the scoping review suggest that QCs improve individual and group performance in terms of costs, ordering of tests, prescription habits or adherence to clinical practice guidelines (Table 1).

**Table 1:**
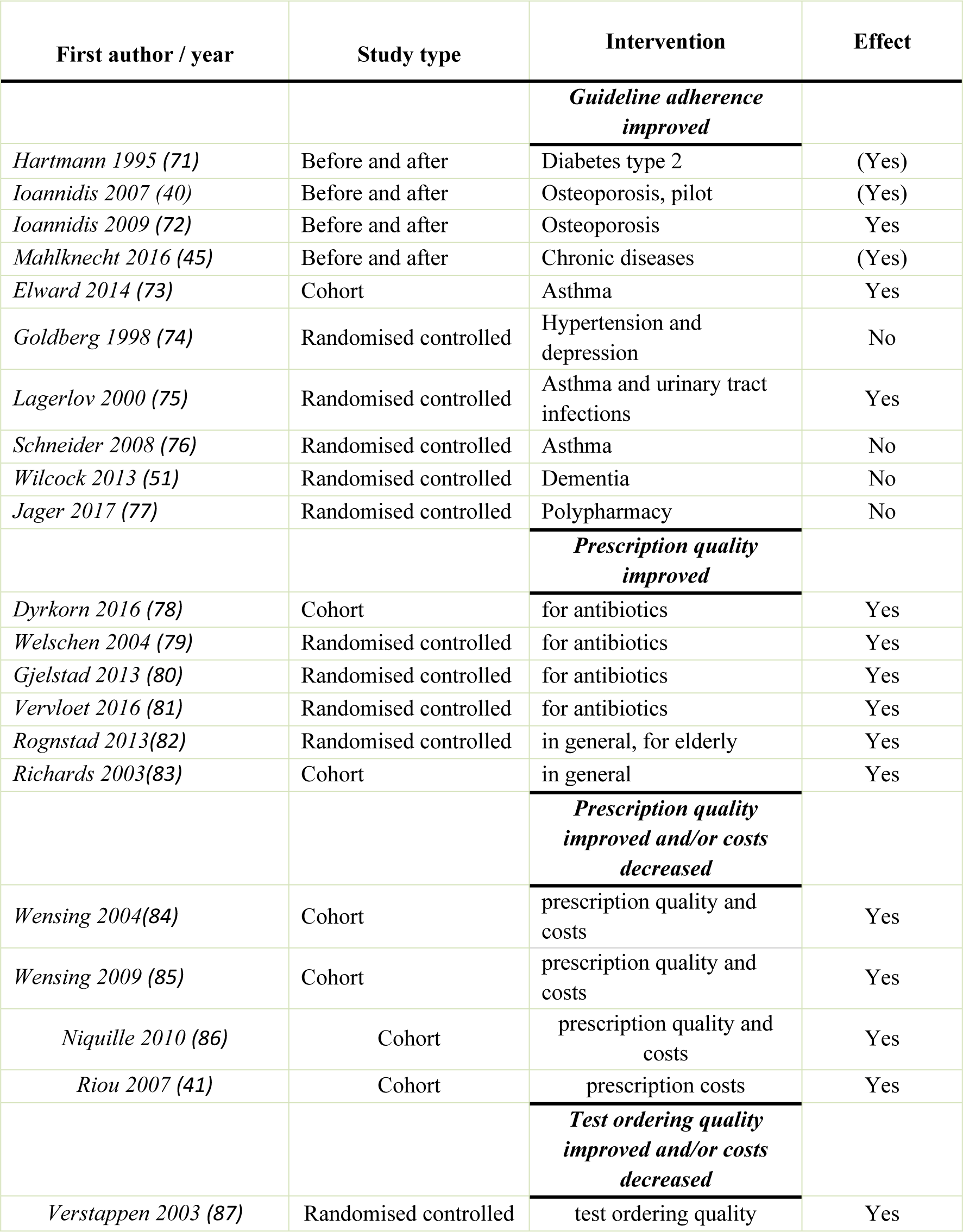
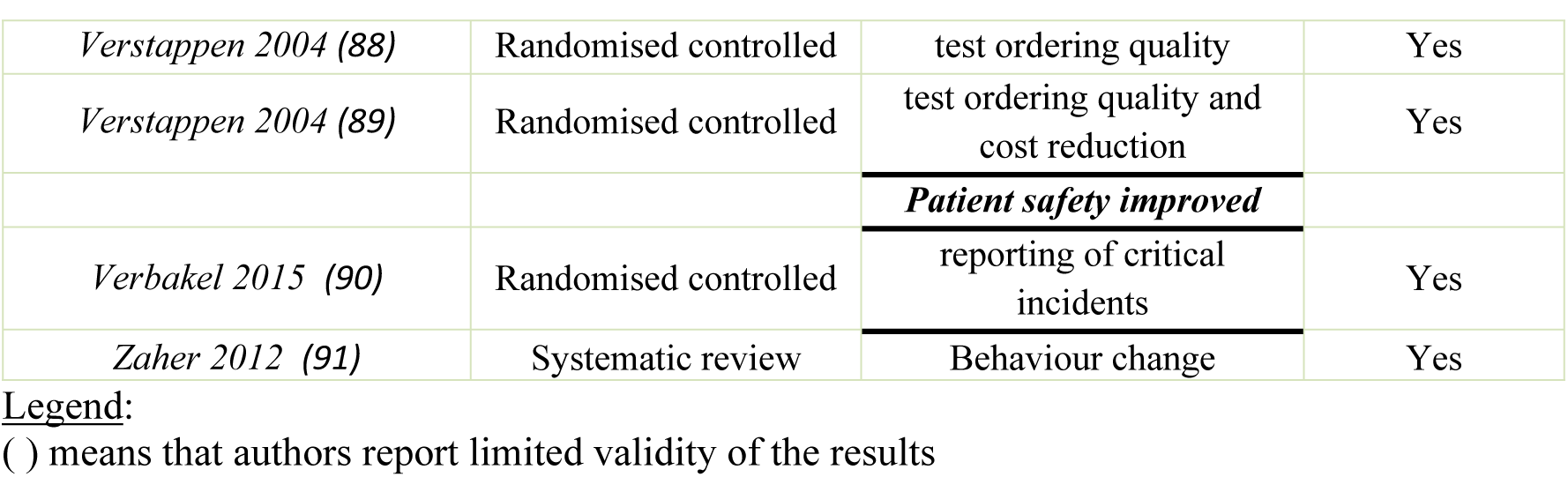
Effectiveness of quality circles

| First author / year | Study type | Intervention | Effect |
| --- | --- | --- | --- |
|  |  | ***Guideline adherence improved*** |  |
| Hartmann 1995 (71) | Before and after | Diabetes type 2 | (Yes) |
| Ioannidis 2007 (40) | Before and after | Osteoporosis, pilot | (Yes) |
| Ioannidis 2009 (72) | Before and after | Osteoporosis | Yes |
| Mahlknecht 2016 (45) | Before and after | Chronic diseases | (Yes) |
| Elward 2014 (73) | Cohort | Asthma | Yes |
| Goldberg 1998 (74) | Randomised controlled | Hypertension and depression | No |
| Lagerlov 2000 (75) | Randomised controlled | Asthma and urinary tract infections | Yes |
| Schneider 2008 (76) | Randomised controlled | Asthma | No |
| Wilcock 2013 (51) | Randomised controlled | Dementia | No |
| Jager 2017 (77) | Randomised controlled | Polypharmacy | No |
|  |  | ***Prescription quality improved*** |  |
| Dyrkorn 2016 (78) | Cohort | for antibiotics | Yes |
| Welschen 2004 (79) | Randomised controlled | for antibiotics | Yes |
| Gjelstad 2013 (80) | Randomised controlled | for antibiotics | Yes |
| Vervloet 2016 (81) | Randomised controlled | for antibiotics | Yes |
| Rognstad 2013(82) | Randomised controlled | in general, for elderly | Yes |
| Richards 2003(83) | Cohort | in general | Yes |
|  |  | ***Prescription quality improved and/or costs decreased*** |  |
| Wensing 2004(84) | Cohort | prescription quality and costs | Yes |
| Wensing 2009 (85) | Cohort | prescription quality and costs | Yes |
| Niquille 2010 (86) | Cohort | prescription quality and costs | Yes |
| Riou 2007 (41) | Cohort | prescription costs | Yes |
|  |  | ***Test ordering quality improved and/or costs decreased*** |  |
| Verstappen 2003 (87) | Randomised controlled | test ordering quality | Yes |
| *Verstappen 2004 (88)* | Randomised controlled | test ordering quality | Yes |
| *Verstappen 2004 (89)* | Randomised controlled | test ordering quality and cost reduction | Yes |
|  |  | ***Patient safety improved*** |  |
| *Verbakel 2015 (90)* | Randomised controlled | reporting of critical incidents | Yes |
| *Zaher 2012 (91)* | Systematic review | Behaviour change | Yes |

20 SRs of high quality and one RCT show that many tools used by QCs predispose professionals to provide care in a different way, enable them to introduce the change, and reinforce it once it has been made (92) (Table 2).

**Table 2:**
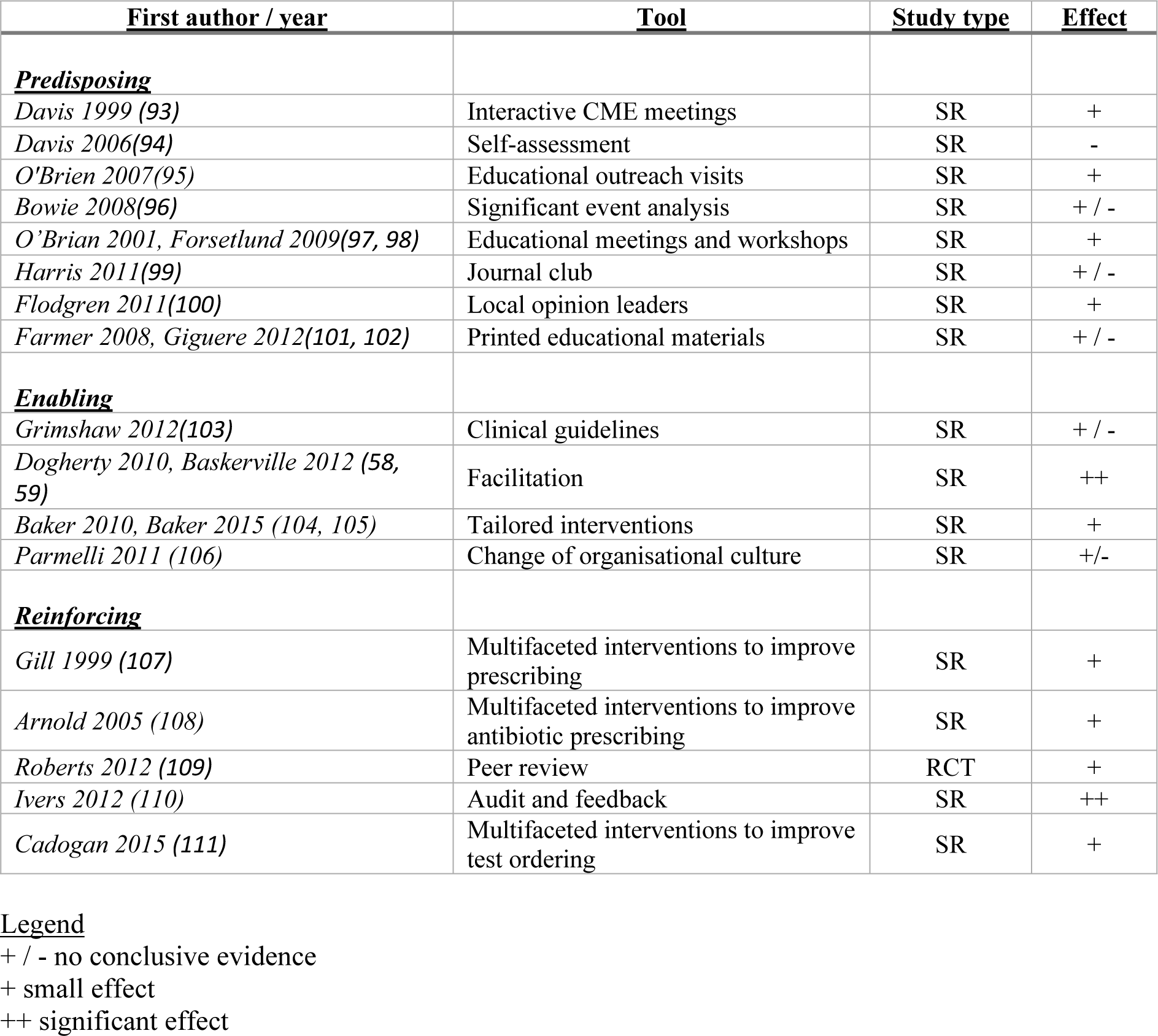
Systematic reviews and randomised controlled trials on tools used in quality circles

| <u>First author / year</u> | <u>Tool</u> | <u>Study type</u> | <u>Effect</u> |
| --- | --- | --- | --- |
| **<u>Predisposing</u>** |  |  |  |
| *Davis 1999 (93)* | Interactive CME meetings | SR | + |
| *Davis 2006(94)* | Self-assessment | SR | - |
| *O'Brien 2007(95)* | Educational outreach visits | SR | + |
| *Bowie 2008(96)* | Significant event analysis | SR | + / - |
| *O'Brian 2001, Forsetlund 2009(97, 98)* | Educational meetings and workshops | SR | + |
| *Harris 2011(99)* | Journal club | SR | + / - |
| *Flodgren 2011(100)* | Local opinion leaders | SR | + |
| *Farmer 2008, Giguere 2012(101, 102)* | Printed educational materials | SR | + / - |
| **<u>Enabling</u>** |  |  |  |
| *Grimshaw 2012(103)* | Clinical guidelines | SR | + / - |
| *Dogherty 2010, Baskerville 2012 (58, 59)* | Facilitation | SR | ++ |
| *Baker 2010, Baker 2015 (104, 105)* | Tailored interventions | SR | + |
| *Parmelli 2011 (106)* | Change of organisational culture | SR | +/- |
| **<u>Reinforcing</u>** |  |  |  |
| *Gill 1999 (107)* | Multifaceted interventions to improve prescribing | SR | + |
| *Arnold 2005 (108)* | Multifaceted interventions to improve antibiotic prescribing | SR | + |
| *Roberts 2012 (109)* | Peer review | RCT | + |
| *Ivers 2012 (110)* | Audit and feedback | SR | ++ |
| *Cadogan 2015 (111)* | Multifaceted interventions to improve test ordering | SR | + |

### Knowledge gaps

All authors of SRs showing effect on behaviour change noted considerable variations within and between studies without being able to account for them. It is difficult to explain in SRs why behaviour change happens in QCs (93). A detailed description of the process of intervention of each step is needed to evaluate how and why they may or may not work (95, 98, 100, 103). This is not only necessary for understanding each step but also for understanding combinations of different interventions or steps, such as the use of printed educational material, combined with the use of local opinion leaders, CME workshop and/or outreach visits (101, 103). It is not known which methods should be used, and under what circumstances, to enable QCs to address the reasons for resisting new practices and barriers to them (105). For example, audit and feedback interventions have typically produced heterogeneous effects, and therefore more exploration is needed to determine the underlying reasons for behaviour change, how best to design and deliver this intervention, when and how to use audit and feedback and, finally, how to optimise this in routine practice (110).

As small group work succeeds in CME, the question arises as to how and why this may or may not work for quality projects as well (54). It seems essential to examine what resources small groups offer GPs for changing behaviour (72). In other words, what it is about QCs that influences the clinical performance of GPs. Further studies are needed to find out how they can be tailored to GPs to achieve better results and what group factors are crucial for better outcomes (85). More information is required to determine how often the group process should be repeated, accepting the fact that once may not be enough (51, 90, 110). As there are hardly any theory-based interventions about change in clinical practice, further research should concentrate on improving our understanding of when, how and why interventions, such as education or providing guidelines, are likely to be effective and how to improve them (111).

### Intentions and benefits of QCs

Knowledge and skills acquired during initial medical education need to be updated through CME, which aims to promote the application of new knowledge via CPD (93, 98, 112, 113). CME and CPD are necessary prerequisites for QI (114–117).

QI is a data-guided activity that brings about positive change in the delivery of care. It deals with local problems like perceived inefficient, harmful or badly-timed health care (118, 119). In some European countries, QCs seem to play a major role in QI, whereas in others they mainly serve CME and CPD (39).

According to qualitative literature, QCs have a number of benefits. Small group work seems to be the preferred way of learning for GPs (47, 53, 116, 120, 121). The groups help them to link evidence to everyday practice (57), learn how to deal with uncertainty (122) and how to improve practice (54). They are a vehicle for discussing issues and reflecting on practice, that may increase self-esteem (123, 124). Frequent participation strengthens team-based strategies for error prevention (125). Participation in groups can mean someone stepping out of their comfort zone when talking about their own practice performance. This may raise anxiety and generate a stress response (124, 126). This same response, however, seems to improve communication skills and provides an opportunity for learning (61, 127). Several groups of authors note that small groups may be an important factor in preventing burnout and for someone remaining in the same area (50, 61, 91, 128–130).

### Origins and Significance of QCs

There are two fundamental concepts that have underpinned the basic understanding of QCs from the beginning of their development: the framework of the Plan-Do-Check-Act Cycle (PDCA) and the social context the group provides for its function (131). In 1924, Shewart created the first table depicting a cycle for continuous control of the QI process (Figure 2) (132). The PDCA cycle is based on the idea that front-line workers often recognize ways of improving production, and will experience increased motivation when given opportunities to participate actively in making those improvements (14, 133). The principles of QI were adopted in health care and presented in three interdependent quality dimensions that interrelate and influence each other: structure, process and outcome. (134).

**Figure 2:**
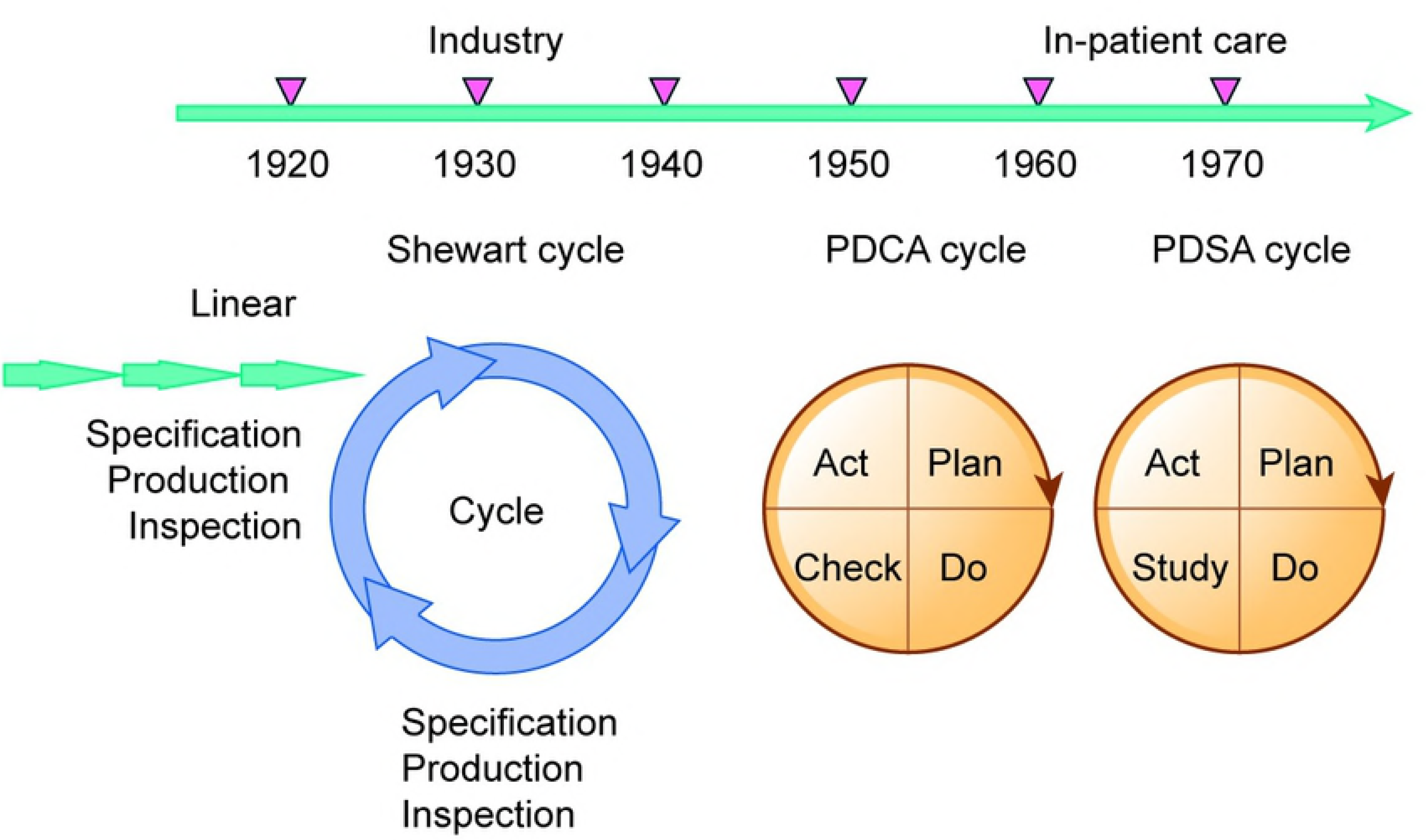
Development of QI Process

Ethical aspects of quality became essential with increasingly discerning patients and public health interests(135) This model of QI in health care was first implemented in in-patient settings and secondary-care clinics in the Netherlands. Development drivers for QCs were the participative group problem-solving approach and the need for shared responsibility for decision-making in fast growing and costly health care systems (136). Ethical aspects of quality became essential with increasingly discerning patients and public health interests

QCs in *PHC* originated in two centres: McMaster University in Canada, and the University of Nijmegen in the Netherlands. Both promoted Problem Based Learning (PBL) where a group of learners are confronted with a problem they have to solve, making them participate actively in gaining knowledge about a particular issue (137).

At McMaster, a practical method using PBL was presented in 1974, whereby GPs met on a regular basis to exchange thoughts about clinical cases and increase and update their knowledge (138). As these groups were primarily concerned with lifelong learning needs, the technique was called Problem Based Small Group Learning (PBSGL) or CME groups.

In 1979, PBL was also implemented experimentally with small groups of GPs in Nijmegen, who met voluntarily on a regular basis, using their peers to continuously and autonomously improve their knowledge (39). As the Netherlands had adopted Donabedian’s dimensions of quality in health care, their small group work contained features of QI. Gradually, the learning cycle transformed into a cycle of QI, as the focus changed from knowledge gain to using knowledge to improve practice (139, 140). PBL added didactic techniques and industrial small group work added communication skills and knowledge about group dynamics.

In subsequent years, the PBSGL method spread from McMaster, Canada, to Ireland, Scotland and England through the building of networks by teachers, academics and policy makers (2, 6–8, 141). Likewise, the European Society for Quality and Safety in Family Medicine (EQuiP) was founded and served as a communication channel for sharing developments, such as QCs, which spread rapidly from the Netherlands to many other European countries, as well as to the USA, Australia and New Zealand, as shown in Figure 3 (1–10, 39, 74, 83, 122, 124, 142–147). In 2015, EQuiP organised a conference in Fischingen, Switzerland, on QCs in PHC where representatives of these very similar movements documented the range of components, characterised their underlying mechanisms and the local context in which they are conducted.

**Figure 3:**
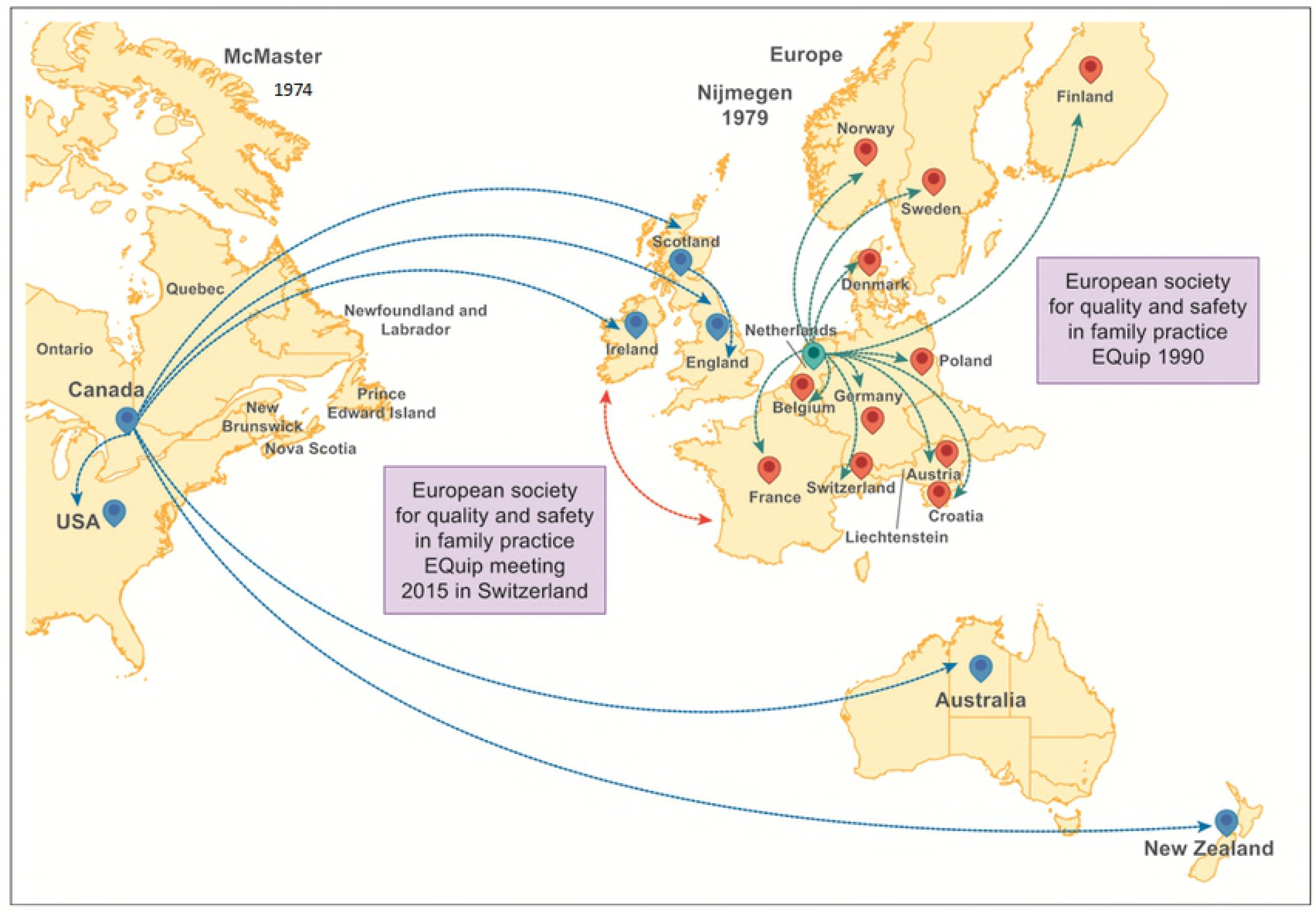
Spread of QCs

## Discussion

### Summary

QCs can change GP behaviour to varying extents and, within the existing SRs and RCTs, authors note small but significant changes in behaviour. Group work appears to fit GP expectations when it comes to CME, CPD and QI projects, where they play a significant role. QCs developed rapidly as the participative group-problem solving approach and the need for shared responsibility became important in societies with spiralling costs for health care.

### Knowledge Gap

The evidence on QCs indicates the existence of substantial knowledge gaps. For example, in studies using methods such as RCT or cohort studies on QCs, or elements thereof, only small and heterogenous effect sizes were noted and it was unclear why these occurred. Further, SRs and RCTs on tools used in QCs have only examined their individual impact or effectiveness. When some have been used in combination, their relative contribution to the overall effect is unclear (107, 108, 111). Finally, it is not known in what way and how many times the process of improvement should be repeated to increase effect sizes(51, 90, 110).

Since QCs embody a complex intervention that is not standardized, and which changes continuously depending on the topic and the context of the group, these results and comments are not surprising (148). Questions regarding the effectiveness of different variations of QC techniques remain unanswered, as well as questions regarding the conditions under which they are most likely to succeed or fail.

### Future Research Directions

Complex interventions such as QCs are difficult to examine. One way of doing this is through the use of realist approaches, namely realist review and realist evaluation. These are theory driven approaches that allow questions to be answered about what works; for whom, how, and why – or not, and in what contexts (148, 149). We have built on the findings of this scoping review and are undertaking a realist review to address these knowledge gaps and research needs (150).

### Strengths and Limitations

To our knowledge, this is the first summary on the origin, significance and effectiveness of QCs in PHC. This review followed accepted methods for undertaking a scoping review and was done in a systematic manner with inbuilt quality assurance processes. Through multiple searches with the input of an expert librarian, we identified a sufficient range of relevant documents that enabled us to fulfil the objectives of the review. Our review is not and was never intended to be a comprehensive summary of evidence regarding QCs. It was designed to clarify working definitions, characteristics and knowledge gaps with a view to planning further research.

## Conclusion

QCs play a major role in CME/CPD and QI. Current evidence indicates that they can be successful but effect size varies substantially. As they are sensitive to local conditions, future research is needed to understand what ingredients and what contextual features lead to successful QCs, using appropriate research techniques such as a realist approach.

## Acknowledgement

Nia Roberts, at the Department of Primary Health Care Sciences, Bodleian Libraries, University of Oxford, has contributed substantially to the search strategy of this work.

## Supporting information

Supplementary file S1: search strings

Supplementary file S2: text books

Supplementary file S3: background papers

Supplementary file S4: key papers on quality circles

